# StackRAM: a cross-species method for identifying RNA N^6^-methyladenosine sites based on stacked ensemble

**DOI:** 10.1101/2020.04.23.058651

**Authors:** Zhaomin Yu, Baoguang Tian, Yaning Liu, Yaqun Zhang, Qin Ma, Bin Yu

**Affiliations:** College of Mathematics and Physics, Qingdao University of Science and Technology, Qingdao 266061, China; Artificial Intelligence and Biomedical Big Data Research Center, Qingdao University of Science and Technology, Qingdao 266061, China; Department of Biomedical Informatics, College of Medicine, The Ohio State University, Columbus, Ohio 43210, USA; School of Life Sciences, University of Science and Technology of China, Hefei 230027, China

**Keywords:** N^6^-methyladenosine sites, Multi-information fusion, Elastic Net, Stacked ensemble method

## Abstract

N^6^-methyladenosine is a prevalent RNA methylation modification, which plays an important role in various biological processes. Accurate identification of the m^6^A sites is fundamental to deeply understand the biological functions and mechanisms of the modification. However, the experimental methods for detecting m^6^A sites are usually time-consuming and expensive, and various computational methods have been developed to identify m^6^A sites in RNA. This paper proposes a novel cross-species computational method StackRAM using machine learning algorithms to identify the m^6^A sites in S*. cerevisiae、*H*. sapiens* and A*. thaliana*. First, the RNA sequences features are extracted through binary encoding, chemical property, nucleotide frequency, k-mer nucleotide frequency, pseudo dinucleotide composition, and position-specific trinucleotide propensity, and the initial feature set is obtained by feature fusion. Secondly, the Elastic Net is used for the first time to filter redundant and noisy information and retain important features for m^6^A sites classification. Finally, the base-classifiers output probabilities are combined with the optimal feature subset corresponding to the Elastic Net, and the combination feature input the second-stage meta-classifier SVM. The jackknife test on training dataset S. *cerevisiae* indicates that the prediction performance of StackRAM is superior to the current state-of-the-art methods. StackRAM prediction accuracy for independent test datasets H. *sapiens* and A. *thaliana* reach 92.30% and 87.06%, respectively. Therefore, StackRAM has development potential in cross-species prediction and can be a useful method for identifying m^6^A sites. The source code and all datasets are available at https://github.com/QUST-AIBBDRC/StackRAM/.

## 1. Introduction

The completion of the Human Genome Project has greatly promoted people’s understanding of the genetic tissue information, transmission, and expression, and made us aware of the extraordinary complexity of the genetic information expression mechanism in cells. As the key link of the central law, RNA connects the genetic material DNA and the performer proteins of life activities. Studies show that there are more than 100 kinds of post-transcriptional RNA modifications, and these modifications are mainly methylated, such as 5-methylcytosine (m^5^C), N^1^-methyladenosine (m^1^A), pseudouridine (ψ), N^6^-methyladenosine (m^6^A) [1,2]. N^6^-methyladenosine (m^6^A) was first found detected in the 1970s, which is one of the most well-studied RNA modifications and widely exists in many species including mammals, plants, bacteria, and viruses [3–6].

As dynamic and reversible process, m^6^A occurs on the sixth nitrogen atom of adenine, and its dynamic change can affect gene expression and cell fate by regulating a variety of RNA-related cell signaling pathways, including mRNA splicing, export, stability, immune tolerance, RNA transcription, processing, cell division, and cell differentiation [7–11]. In addition, m^6^A modification is closely related to human diseases, such as cancer [12], virus infection [13], and brain development abnormalities [14]. Therefore, accurate identification of the m^6^A sites is essential for basic research on RNA methylation modification, and also important for drug development, understanding disease mechanisms and promoting the bioinformatics development. The experimental methods to identify m^6^A sites in RNA sequences are two-dimensional thin layer chromatography [15], high performance liquid chromatography [16], and high-throughput methods (m^6^A-seq [17] and MeRIP-Seq [14]). The biochemical experimental approaches for detecting m^6^A sites are costly and time-consuming, and identifying exact positions of m^6^A sites in RNA is still challenging [18]. With the development of advanced sequencing technology and genome projects, a large number of RNA sequences have been accumulated, and many researchers have proposed effective computational methods based on machine learning algorithms for fast and accurate prediction of m^6^A sites.

Until now, researchers developed a series of m^6^A sites computational methods based on machine learning. Huang et al. [19] proposed a cross-species classifier BERMP to predict the m^6^A sites by integrating a deep learning algorithm and a random forest approach. Zhao et al. [20] constructed the model HMpre to resolve the imbalance data issues in the human mRNA m^6^A prediction problem with the cost-sensitive approach. HMpre on independent test dataset achieves a much better performance of precision 0.3035, F1 0.3961 and MCC 0.3329. Chen et al. [21] constructed iRNA-PseDNC to identify m^6^A sites using pseudo nucleotide composition. It has been demonstrated by the 10-fold cross-validation that the performance of iRNA-PseDNC is superior to RAM-NPPS. Chen et al. [22] developed the computational method RAM-ESVM, which employed ensemble support vector machine classifiers to predict the N^6^-methyladenosine sites in the RNA. The jackknife test results showed that RAM-ESVM outperforms a single support vector machine. Wang et al. [23] proposed a new tool RFAthm^6^A predicting m^6^A sites in *Arabidopsis thaliana*. Akbar et al. [24] constructed model iMethyl-STTNC model based on SVM, which by extending the idea of SAAC into Chou’s PseAAC to predict N^6^-methyladenosine sites. SVM using STTNC feature space reported accuracy of 69.84%, 91.84% on dataset1 and dataset2, respectively. Zhang et al. [25] developed a computational method for identifying RNA m^6^A sites in *Escherichia coli* genome and the accuracy values obtained by the proposed method are more than 90% in both 10-fold cross-validation test and independent dataset test. Zhang et al. [26] constructed the predictor m^6^A-HPCS based on the HPCS algorithm, which can effectively extract optimized subsets of nucleotide physical-chemical properties. Xiang et al. [27] constructed a model RNAMethPre based on the support vector machine to predict the m^6^A site in mRNA. Qiang et al. [28] proposed a prediction model m^6^AMRFS based on eXtreme Gradient Boosting (XGBoost), which uses dinucleotide binary encoding and local position-specific dinucleotide frequency to extract RNA sequences. Chen et al. [29] proposed a prediction tool MethyRNA for identification of N^6^-methyladenosine sites based on SVM, and the accuracy of H*. sapiens* and M*. musculus* reached 90.38% and 88.39%, respectively.

Although researchers have made great contributions to RNA methylation modification studies, and most of the models for m^6^A sites prediction are single traditional classifier. Inspired by this, this paper proposes a novel method StackRAM for identifying m^6^A sites based on the stacked ensemble method and uses the most fairly and strictly jackknife test to evaluate the model prediction performance. First, we select binary encoding, chemical property, nucleotide frequency, k-mer nucleotide frequency, pseudo dinucleotide composition and position-specific trinucleotide propensity to extract the RNA sequence features of different species, and convert the sequence information into numerical information recognized by machine learning algorithms. Secondly, compared with the other six dimensional reduction methods, the Elastic Net is selected as the best dimensional reduction method, which is performed on the initial feature space fused with multiple features. Then, the optimal feature subset corresponding to the Elastic Net is input into AdaBoost, ERT, KNN, XGBoost, RF, LightGBM, and SVM to identify the m^6^A sites, respectively. By comparing the m^6^A sites prediction accuracy of different classifiers, LightGBM and SVM are selected as the best base-classifiers. In addition, the prediction probability values of the first stage in StackRAM and the optimal feature subset are combined as the input features of the second stage. The prediction accuracy of combined features is compared between LR and SVM, and SVM is selected as the meta-classifier. Finally, the jackknife test is used to examine prediction method effectiveness, and StackRAM is compared with other state-of-the-art prediction models. At the same time, independent test datasets are used to verify the generalization performance of the prediction method proposed in this paper, and the results show that the StackRAM is more competitive in m^6^A sites prediction.

## 2. Materials and methods

### 2.1 Datasets

High-quality datasets are the foundation for constructing and training accurate and effective prediction models. In this paper, we select real datasets of *Saccharomyces cerevisiae* (S. *cerevisiae*)*, Homo sapiens* (H. *sapiens*) and *Arabidopsis thaliana* (A. *thaliana*) to evaluate the prediction performance of the proposed model, and the datasets contain both positive samples and negative samples. Dataset S. *cerevisiae* is used as the training dataset, and datasets H. *sapiens* and A. *thaliana* are used as the independent test datasets. The S. *cerevisiae* dataset is downloaded from Chen’s work [30], and all the sequence samples are 51-nt long with the consensus motif GAC. If the sample sequence does not have nucleotides at certain positions, the missed nucleotides will fill in its mirror image. The dataset includes 1307 positive sequences, to avoid the imbalanced dataset impact on the robust model construction, and the equal number of negative sequences are picked out from a negative dataset which contains 33280 samples. The H. *sapiens* dataset is downloaded from Feng’s work [31], and the window size of samples is 41. If the length of the sequence sample is less than the window size, the lacking nucleotide is filled with the same nucleotide of its nearest neighbor. In order to avoid generating highly skewed datasets, the sample numbers of the positive and negative datasets are kept consistent, both of which are 1130. The A. *thaliana* dataset is derived from the research of Chen et al. [32], using the CD-HIT to reduce sequence homology bias and remove sequences with more than 60% sequence similarity. The dataset contains 394 positive samples and 394 negative samples, and all the samples are with the length of 25 nucleotides.

### 2.2 Feature encoding algorithm

In theory, the nucleotides of RNA sequences contain all the necessary information for the m^6^A sites prediction. In order to encode the RNA sequence into a vector recognized by machine learning, this paper uses binary encoding [33], chemical property (NCP) [34], nucleotide frequency (ANF) [35], k-mer nucleotide frequency (K-mer) [36], pseudo dinucleotide composition (PseDNC) [37] and position-specific trinucleotide propensity (PSTNP) [38] to extract RNA sequence features.

#### 2.2.1 Binary encoding

Binary encoding is a common feature extraction method to characterize RNA sequences and can accurately describe the nucleotides at each position in the sample sequence. Since the RNA sequence contains four nucleotides adenine (A), guanine (G), cytosine (C), and uracil (U), this feature extraction method encodes each nucleotide into a 4-dimensional binary vector of (1, 0, 0, 0), (0,1, 0, 0), (0, 0,1, 0) and (0, 0, 0,1), respectively. Therefore, the binary encoding of *L* -nt window will result in a 4 × *L* dimensional feature vector.

#### 2.2.2 Chemical property

RNA consists of four types of nucleotides: A, C, G, and U. Based on differences in chemical property, they can be categorized into three different groups. For the number of rings, adenine and guanine have two rings, and cytosine and uracil have one ring. In terms of chemical functionality, adenine and cytosine contain amino groups, while guanine and uracil contain keto groups. For secondary structures formation, guanine and cytosine have strong hydrogen bonds, while adenine and uracil have weak hydrogen bonds. Therefore, each nucleotide in the RNA sequence is encoded into vector *s*_*i*_ (*x*_*i*_, *y*_*i*_, *z*_*i*_) according to formula (1).

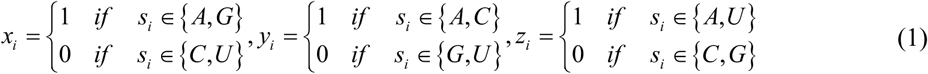

According to the above three division methods, ‘*A’* can be represented by vector (1,1,1), ‘*C ‘* can be represented by vector (0,1, 0), ‘*G ‘* can be formulated as vector (1, 0, 0), and ‘*U ‘* can be formulated as vector (0, 0,1). Therefore, *L* -nt long RNA sequence could be generated a 3 × *L* -dimensional feature vector.

#### 2.2.3 Nucleotide frequency

In order to understand the composition and frequency of nucleotides near the m^6^A sites, the density *d*_*i*_ of nucleotide *n*_*j*_ at the RNA sequence specific position *i* is calculated as follows:

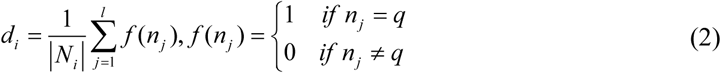

where *N*_*i*_ denotes the length of the *i* -th subsequence and *l* denotes the nucleotide position in the subsequence (*q* ∈ {*A*, *C*, *G*,*U*}). For instance, suppose the RNA sequence is ‘*GAUCACCG*’. The density of ‘*A*’is 1/2, 2/5 at sequence positions 2 and 5, respectively. The density of ‘*C*’at the sequence position 4, 6, or 7 is 1/4, 1/3, or 3/7, respectively. The density of ‘*G*’is 1, 1/4 at sequence positions 1 and 8, respectively. The density of ‘*U*’is 1/3 at sequence position 3. When we calculate the density of each position in sample sequence, the occurrence density at the first position is always 1. Therefore, sequence with length *L* is represented by *L* − 1 dimensional feature vector.

#### 2.2.4 K-mer nucleotide frequency

Adjacent nucleotide pairs affect the structure and function of the RNA sequence, reflecting the sequence background differences between the m^6^A sites and the non-m^6^A sites. The K-mer nucleotide frequency is used to calculate the frequency of adjacent nucleotides in the sample sequence, 4^*K*^-dimensional feature vector will be generated. As K increases, the dimension of the feature vector increases exponentially, leading to overfitting problems in the prediction model. Therefore, the 2-mer nucleotide frequency is employed to encode the sample sequence, and calculate the frequency of *AA*, *AC*, *AG*, *AU*, *CA*, *CC*, *CG*, *CU*, *GA*, *GC*, *GG*, *GU*, *UA*,*UC*,*UG*,*UU* in the RNA sequence to generate 16-dimensional feature vector.

#### 2.2.5 Pseudo dinucleotide composition

In order to fuse the local and global sequence information of the RNA sequence, we use pseudo dinucleotide composition to extract the feature information of the RNA nucleotide. Each sample sequence will generate a feature vector as shown in formula (3):

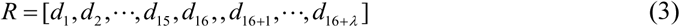

Among them:

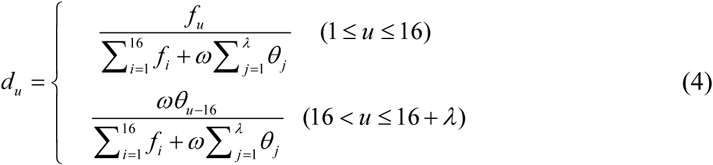

In Eq. (4), the first 16 components are used to incorporate short-range or local sequence pattern information in the RNA sequence, and the remaining elements represent long-range or global sequence pattern information. *λ* is the total pseudo component to reflect long-range or global sequence effect, *ω* is the weight factor, *f*_*u*_ is the normalized frequency of non-overlapping dinucleotides at the *u*-th position in the RNA sequence, and *θ*_*j*_ is the *j*-th tier correlation factor.

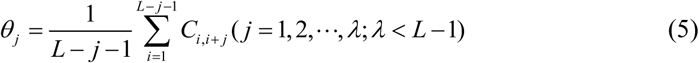

where, *θ*_1_ is the first-tier correlation factor, which reflects the sequence order correlation of all the dinucleotides that are most adjacent along the RNA sequences, *θ*_*j*_ is *j*-th tier correlation factor that reflects the sequence order correlation between all the *j*-th most adjacent dinucleotides in the RNA sequences. The coupling factor *C*_*i,i+j*_ as described by the following formula (5):

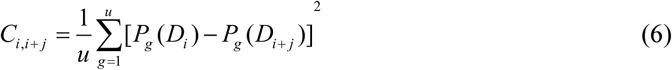

where *u* is the number of physicochemical properties of RNA and *P*_*g*_ (*D*_*i*_) normalized and defined as follows:

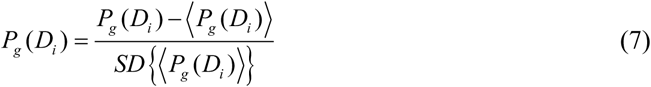

where the symbol 〈〉 represents the average of the quantity of the different dinucleotides and *SD* means the corresponding standard deviation.

#### 2.2.6 Position-specific trinucleotide propensity

Single-strand position-specific trinucleotide propensity (PSTNP) is used to describe the statistical significance of RNA. For RNA sequences, 4^3^ = 64 trinucleotides will generate: *AAA*, *AAC*, *AAG*,…,*UUU*. Therefore, for an RNA sequence sample with length *L*, the trinucleotide position-specific can be expressed by the matrix of 64×(*L* − 2):

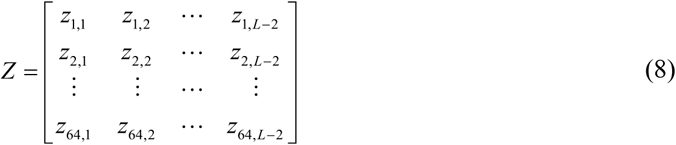

where *z*_*i,j*_ = *F*^+^(3*mer*_*i*_|*j*)-*F*^−^(3*mer*_*i*_|*j*) and (*i* = 1,2,…, 64; *j* = 1,2,… *L* − 2)

*F*^+^(3*mer*_*i*_|*j*) and *F*^−^(3*mer*_*i*_|*j*) represent the frequency of occurrence of the *i*-th trinucleotide at the *j*-th position in the positive dataset *S* and the negative dataset *S*, respectively. 3*mer*_1_ denotes *AAA*, 3*mer*_2_ denotes *AAC*, …, 3*mer*_64_ denotes *UUU*.

For a given RNA sample sequence, it can be expressed as:

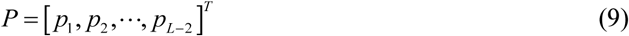

where *T* is the transpose operator and *p*_*u*_ is defined as follows:

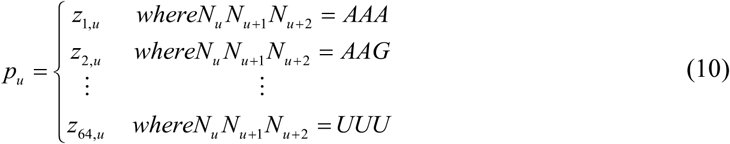

### 2.3 Elastic Net

In order to reduce the model overfitting risk, Tibshirani [39] introduced *ℓ*_1_ norm regularization and proposed the least absolute shrinkage and selection operator (Lasso) in 1996. However, the Lasso method does not apply to the case where there is some connection in the feature space, that is, there are multiple relevant features. Therefore, Zou and Hastie [40] proposed the Elastic Net method based on Lasso theory, using *ℓ*_1_ and *ℓ*_2_ norm regularization for training. This combination allows learning sparse models, when faced with relevant features, Lasso may choose one randomly, while Elastic Net may choose both.

Elastic Net objective function is defined as follows:

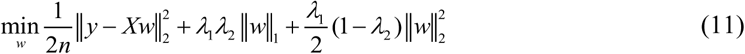

where, *λ*_1_ and *λ*_2_ are non-negative penalty parameters, and *λ*_2_ represents the scaling between the *ℓ*_1_ and *ℓ*_2_ penalty in Elastic Net, which ranges from *λ*_2_ = 0 to 1. For *λ*_2_ = 0, the penalty is *ℓ*_2_, which is Ridge. For *λ*_2_ = 1, the penalty is *ℓ*_1_, which is Lasso. For 0 < *λ*_2_ < 1, the penalty is a combination of *ℓ*_1_ and *ℓ*_2_.

### 2.4 Framework of StackRAM

Ensemble learning is the model based on multiple classifiers and uses a certain rule to integrate a series of learning results to obtain better results than the single classifier. This paper builds StackRAM based on the stacked ensemble method to identify RNA methylation modification sites. The stacked ensemble minimizes the generalization error by combining the information from multiple prediction models, and it is applied in the bioinformatics field [41–44]. It mainly consists of two stages of learning, including the first stage base-classifier and the second stage meta-classifier. The first stage train base-classifier based on the initial dataset and the second stage uses the probability output of the base-classifier as input features to correct inaccurate training at the first stage and reduce generalization errors for training second stage meta-classifier.

Different classifiers have different feature learning spaces and class recognition capabilities. In the first stage, we explore 7 different machine learning algorithms to select the base-classifier of the stacked ensemble method, including AdaBoost, ERT, KNN, XGBoost, RF, LightGBM and SVM. By comparing the prediction accuracy of 7 machine learning algorithms on the optimal feature subset, two LightGBMs and two SVMs are used as base-classifiers. The meta-classifier integrates the probability output values of multiple base-classifier in the first stage and learns the relationship between different predictors and real class labels to enhance the model’s prediction performance. In the second stage, we take the optimal feature subset and the probability output of the first stage as new combined features, which are input to SVM and LR, respectively, and choose SVM as the meta-classifier according to the prediction results. StackRAM can mine the essential abstract features that characterize RNA methylation sites through hierarchical learning, and its prediction performance is better than that of the single classifier. The stacked ensemble algorithm is described in Algorithm 1.

#### Algorithm 1 Stacked ensemble algorithm

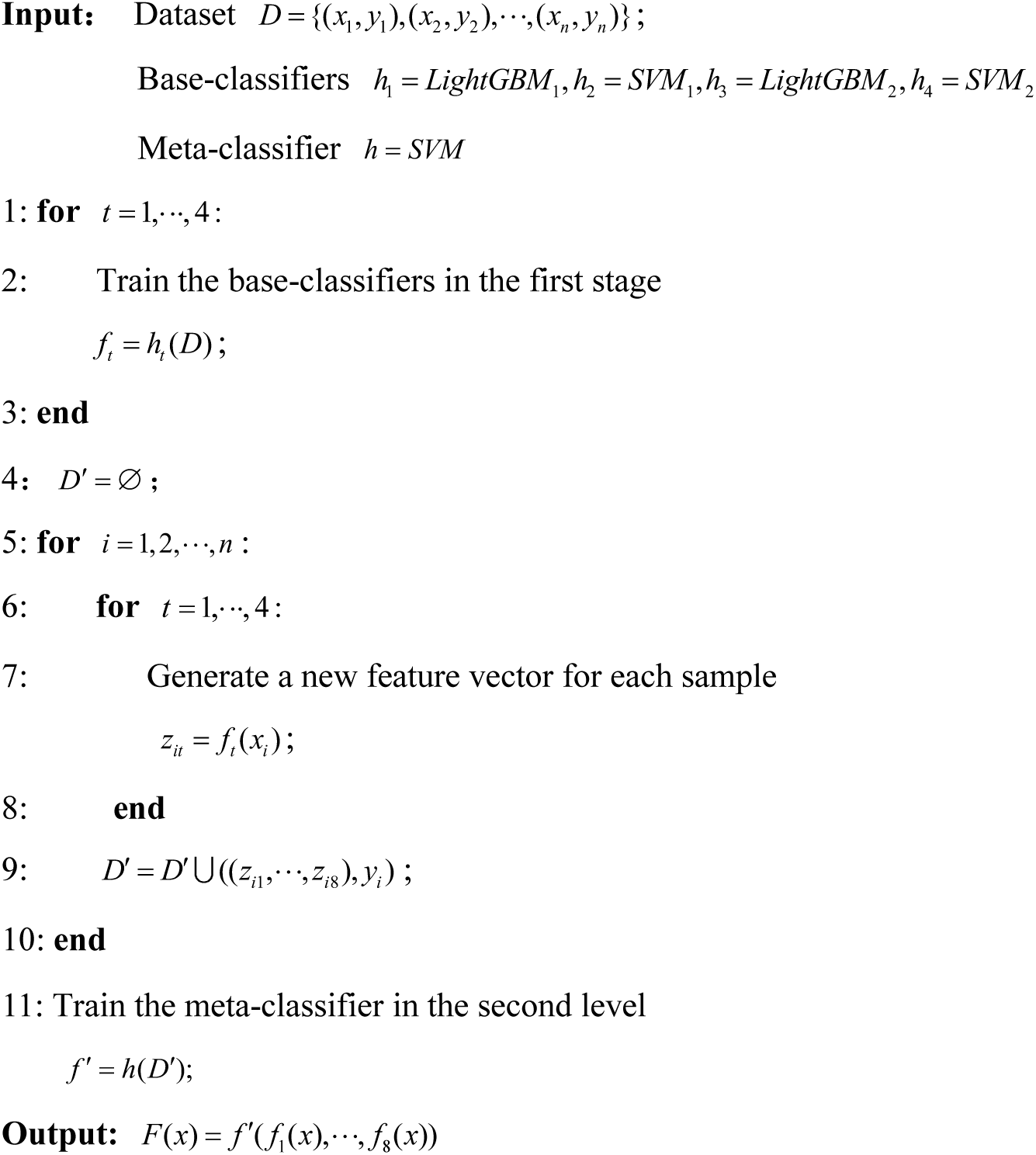

### 2.5 Performance evaluation

In statistical prediction, the following three methods are used to evaluate the effectiveness of the model: independent dataset test [45], K-fold cross-validation [46], and jackknife test [47]. As the most objective and involuntary cross-validation method, the jackknife test is adopted to examine the predictive quality of a computational model by more and more researchers. In this paper, the most accurate and rigorous jackknife test is selected to evaluate model performance. Each sample in the dataset is in turn singled out as an independent test dataset, and the remaining samples are used as the training dataset to train the model. In order to examine the quality of the prediction model fairly and objectively, four common metrics are employed: sensitivity (Sn), specificity (Sp), accuracy (ACC), and Mathew’s correlation coefficient (MCC), which are defined as:

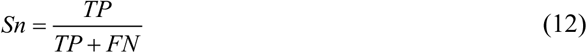

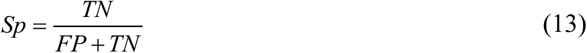

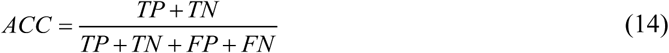

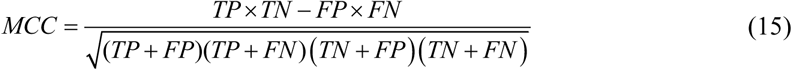

where, *TP TN FP FN* respectively denote the number of true positives, true negatives, false positives, and false negatives. *TP* indicates the number of true m^6^A sites that are recognized as m^6^A sequences correctly, *TN* indicates the number of non-m^6^A sites that are recognized as non-m^6^A sequences correctly, *FP* represents the number of non-m^6^A sites that are predicted true m^6^A sites, and *FN* represents the number of m^6^A sites that are predicted true non-m^6^A sites. Sn and Sp represent the model’s ability to correctly predict positive and negative samples, while ACC and MCC are used to evaluate the overall performance of the model. In addition, ROC curves [48] and PR curves [49] are also used to evaluate the robustness and prediction performance of the model. The ROC curves plot the relationship between the sensitivity and 1-specificity, and the precision-recall (PR) curves plot the relationship between precision and recall (sensitivity). The area values under the ROC curve and PR curve are AUC and AUPR, respectively. The AUC and AUPR near 1 imply perfect prediction performance.

### 2.5 The StackRAMs pipeline

The m^6^A sites prediction method proposed in this paper called StackRAM and the calculation process is shown in Fig. 1. All the experiments are carried out on a Windows Server 2012R2 Intel (R) Xeon (TM) CPU E5-2650 @ 2.30GHz 2.30GHz with 32.0GB of RAM, MATLAB2014a and Python 3.6 programming implementation.

**Fig. 1.**
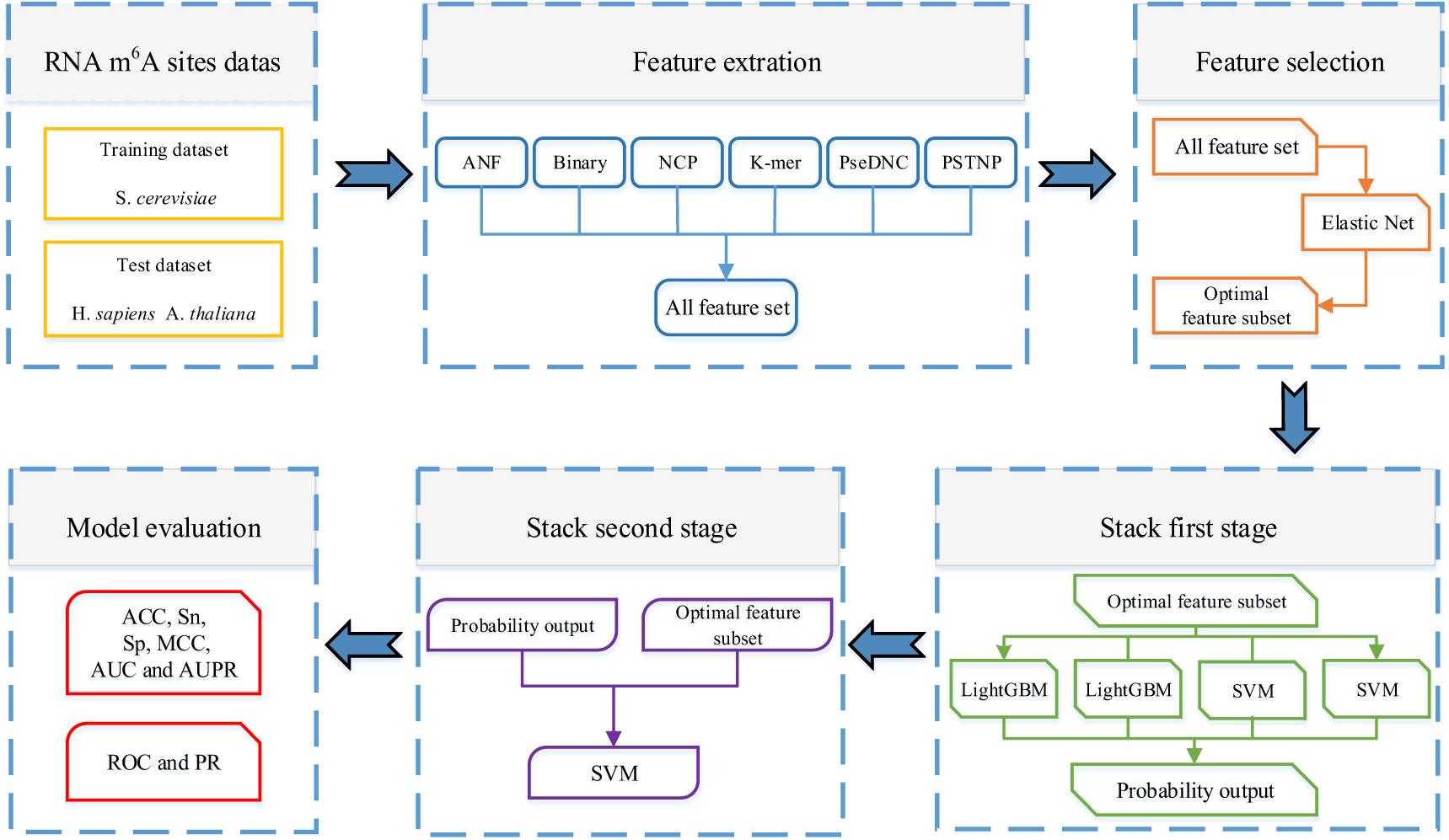
The pipeline of the StackRAM method **f**or identifying RNA N^6^-methyladenosine sites.

The detailed steps of the StackRAM model are described as follows:

***Step 1:*** Obtain RNA methylation modification datasets of three different species, including the RNA positive and negative sample sequences of the datasets and the corresponding class labels.

***Step 2:*** Feature encoding. Use the sequence-derived information to encode the RNA sequence, convert the character information into numerical vectors, determine the optimal parameter of PseDNC, fuse the feature vectors containing 6 different types of information, and obtain the initial feature space of the original dataset.

***Step 3:*** Feature selection. For the initial feature space, Elastic Net is used to reduce the dimension, remove redundant and noisy features, retain important features related to model classification, and get the optimal feature subset.

***Step 4:*** Classification algorithm. In the first stage, for the optimal feature subset, two LightGBM and two SVM are selected as base-classifiers to identify whether the input RNA sequence contains m^6^A sites. In the second stage, SVM is selected as the meta-classifier for the combined features of the probability output of the first stage and the optimal feature subset.

***Step 5:*** According to ***Step 2*** to ***Step 4***, input the feature dataset of the training dataset and the corresponding category labels into StackRAM to predict the m^6^A sites of the dataset S. *cerevisiae. **Step 6:*** Model performance evaluation. The m^6^A sites of H. *sapiens* and A. *thaliana* are identified according to the training model, and Sn, Sp, MCC, ACC, AUC, and AUPR values are calculated using the jackknife method, and ROC and PR curves are drawn to evaluate the predictive performance of the model.

## 3. Results and discussion

### 3.1 Nucleotide composition analysis

In order to analyze the difference in nucleotide distribution between the m^6^A sites and the non-m^6^A sites in the RNA sequence, identify the nucleotide information near the m^6^A sites and reveal the nucleotide pattern around the m^6^A sites. In this paper, Two-Sample Logos [50] (http://www.twosamplelogo.org/cgi-bin/tsl/tsl.cgi) is used to determine and visualize the statistically significant differences between the positives and negatives of the training dataset S. *cerevisiae* as shown in Fig. 2.

**Fig. 2.**
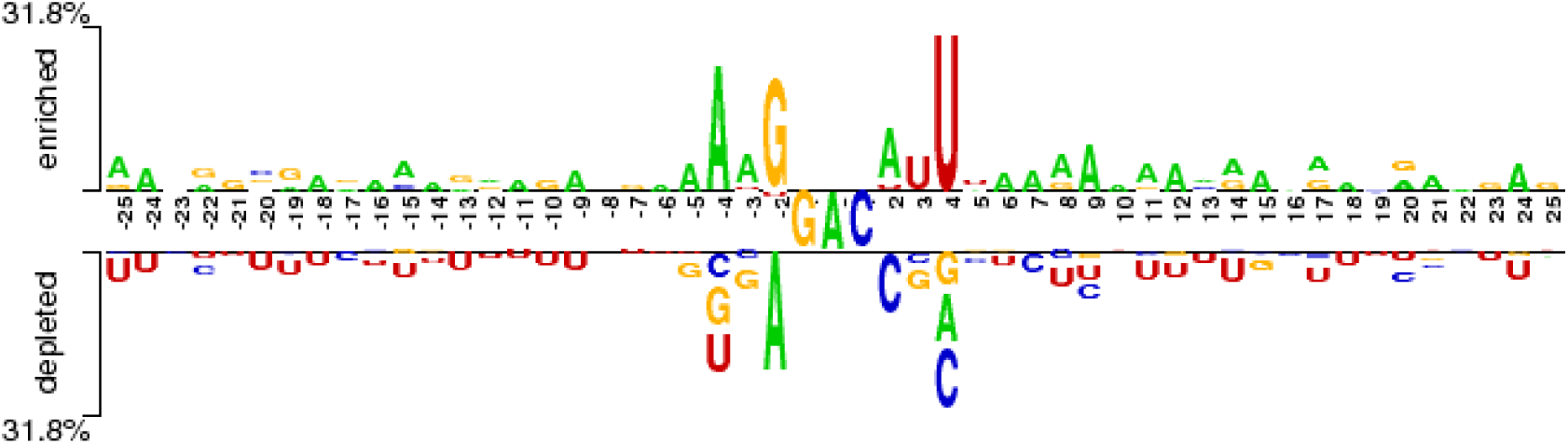
The nucleotide composition preference of sequences between m^6^A and non-m^6^A sites in S. *cerevisiae*. The top vertical axis is the composition preference of m^6^A sites containing sequences, while the down vertical axis is the composition preference of non-m^6^A sites containing sequences.

As shown in Fig. 2, there are significant differences between the nucleotide composition near the m^6^A sites and the non-m^6^A sites in the RNA sequence of S. *cerevisiae*. Dataset S. *cerevisiae* near the m^6^A sites, adenine occurrence frequency is higher at positions −4, −3, and 2, guanine occurrence frequency is higher at positions −2, −1, uracil occurrence frequency is higher at positions 3 and 4, and cytosine occurs more frequently at position 1. Dataset S. *cerevisiae* near non-m^6^A sites, cytosine occurs more frequently at positions 2 and 4, and adenine occurs more frequently at position −2. Based on these characteristics, there are different compositional and positional information between m^6^A sites and the non-m^6^A sites in S. *cerevisiae*.

### 3.2 Impact of the parameter *λ* the PseDNC

In the PseDNC feature extraction method,. *λ* reflects global sequential pattern effect. The larger the value, the more global sequential pattern information of the model is included. In order to explore the influence of the parameter *λ* on the PseDNC feature extraction method, and considering the dataset minimum window length is 25, we set *λ* values to 4, 8, 12, 16, 20, and 23, respectively. The base-classifiers LightGBM and SVM are selected to identify the m^6^A sites of the training dataset S. *cerevisiae*, and the prediction accuracy values of the training dataset to different parameters *λ* are shown in Fig. 3.

**Fig. 3.**
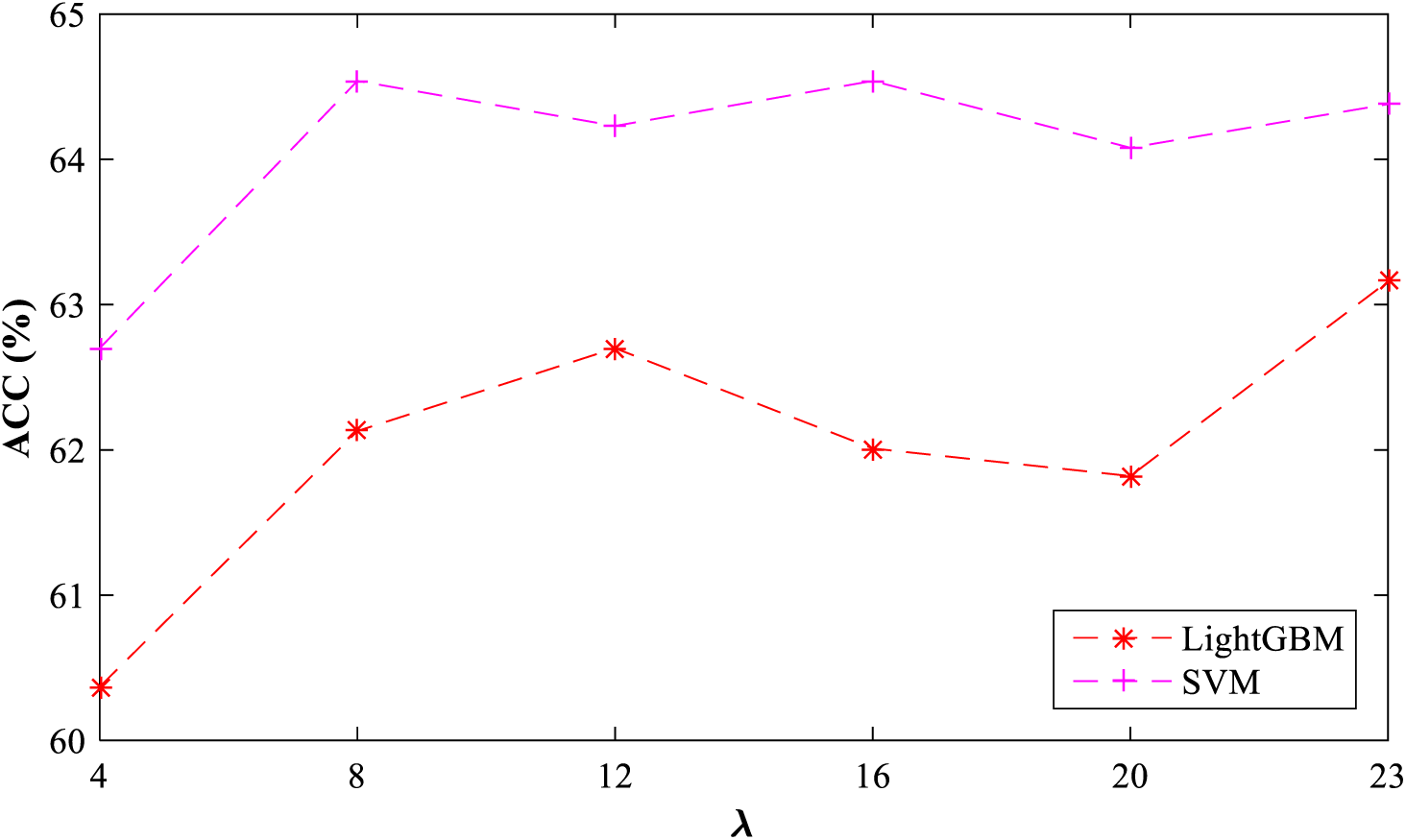
ACC values to different parameters *λ* in PseDNC on S. *cerevisiae*.

From Fig. 3 we can intuitively see that for the classifiers LightGBM and SVM, the dataset S. *cerevisiae* has different prediction accuracy values for different parameters in PseDNC. For the base-classifier LightGBM, the prediction accuracy of the dataset S. *cerevisiae* to the parameter *λ* is first increased and then decreased. When the *λ* value is 23, the prediction accuracy reaches the highest value, which is 63.16% (Supplementary Table S1). For the base-classifier SVM, the prediction accuracy is first increased and then relatively stable. When the parameter *λ* is 8, and prediction accuracy ACC reaches the maximum value, which is 64.54%. When the parameter *λ* is 23, the prediction accuracy achieves 64.38% (Supplementary Table S1). Considering the influence of the parameter *λ* on the two base-classifiers, the optimal parameter *λ* in PseDNC is set to 23, and a 39-dimensional feature vector is generated for each sample sequence.

### 3.3 Comparative analysis of different feature extraction methods

One of the challenges in accurately identifying m^6^A sites is extracting information-rich features. Because a single feature extraction method is not sufficient to obtain the difference between m^6^A sites and non-m^6^A sites, this paper uses six sequence-derived feature extraction methods to extract RNA nucleotide sequence information. The RNA sequences are converted into high discrimination and high-quality feature vectors, which improve the model’s ability to accurately predict m^6^A sites. In order to compare the effects of different feature extraction methods on m^6^A sites prediction, nucleotide frequency (ANF), binary encoding (Binary), chemical property (NCP), k-mer nucleotide frequency (K-mer), pseudo dinucleotide composition (PseDNC), position-specific trinucleotide propensity (PSTNP) corresponded the single feature dataset and the fused feature dataset All are input to the base-classifiers LightGBM and SVM, respectively. The accuracy values of different feature extraction methods on the training dataset S. *cerevisiae* m^6^A sites are shown in Table 1, and the corresponding dimensions of different feature extraction methods are shown in Supplementary Table S2.

**Table 1.** Prediction accuracy values of training dataset S. *cerevisiae* on different feature extraction methods.

| ACC (%) | ANF | Binary | NCP | K-mer | PseDNC | PSTNP | All |
| --- | --- | --- | --- | --- | --- | --- | --- |
| LightGBM | 61.29 | 72.07 | 72.72 | 61.59 | 63.16 | 79.42 | 80.15 |
| SVM | 61.55 | 72.88 | 73.22 | 63.12 | 64.38 | 80.72 | 78.88 |

It can be clearly seen from Table 1, the classifiers LightGBM and SVM have different prediction accuracy for different feature extraction methods of S. *cerevisiae*, that is, different feature extraction methods have different contributions to m^6^A site recognition. For the single feature extraction method PSTNP, the highest prediction accuracy of the base-classifiers LightGBM and SVM are highest, which are 79.42% and 80.72%, respectively. The ACC values corresponding to the base-classifiers LightGBM and SVM are 18.13% and 19.17% higher than the values corresponding to feature extraction method ANF, 17.83% and 17.60% higher than the values corresponding to feature extraction method K-mer. The feature extraction methods Binary and NCP have higher prediction accuracy. The prediction accuracy of the classifier LightGBM reaches 72.07% and 72.72%, and the prediction accuracy of the classifier SVM reaches 72.88% and 73.22%. The accuracy values of PseDNC’s base-classifiers on the dataset S. *cerevisiae* reach 63.16% and 64.38%, respectively, indicating that it can fuse local and global sequence information of RNA sequences.

The prediction accuracy of the base-classifier LightGBM on the fused feature dataset All is higher than 6 single feature extraction methods, indicating that multi-information fusion can integrate multiple types of information and improve the model’s prediction accuracy to a certain extent. The prediction accuracy of the base-classifier SVM for All is only lower than the corresponding value of PSTNP, which indicates that multiple information fusion will produce redundant features and reduce the model prediction accuracy.

### 3.4 Comparative analysis of different feature selection methods

Although multi-information fusion improves the prediction accuracy of the model to a certain extent, it also brings redundant feature information, which affects the prediction bias of the model. Besides, high-dimensional features will affect the calculation speed of the model. In this section, we perform feature optimization on the fused feature set All to determine the best dimensional reduction method. In this paper, the jackknife method is used on S. *cerevisiae*. Locally linear embedding (LLE) [51], max-relevance-max-distance (MRMD) [52], spectral embedding (SE) [53], singular value decomposition (SVD) [54], mutual information (MI) [55], extra-trees (ET) [56] and Elastic Net are selected for feature selection of the fused feature space. The Elastic Net penalty parameter *λ*_1_ is set to 0.1, and the penalty parameter *λ1*_2_ is set t 0.05, which eliminates 346-dimensional redundant features in the original feature dataset and retains 165-dimensional features with important significance for model identification (Supplementary Table S2 and Table S3). In order to better compare with Elastic Net, the feature subsets corresponding to the dimensional reduction methods of LLE, MRMD, SE, SVD, MI on training dataset S. *cerevisiae* are all 165 dimensions. The feature subset corresponding to the dimensional reduction method ET contains 259-dimensional features. By inputting the feature subsets corresponding to different dimensional reduction methods into the base-classifiers LightGBM and SVM, respectively, the accuracy values of the training dataset S. *cerevisiae* for different dimensional reduction methods are shown in Table 2.

**Table 2.** Prediction accuracy of training dataset S. *cerevisiae* on different dimensional reduction methods.

| ACC (%) | LLE | MRMD | SE | SVD | MI | ET | Elastic Net |
| --- | --- | --- | --- | --- | --- | --- | --- |
| LightGBM | 69.01 | 73.11 | 75.13 | 75.86 | 77.31 | 79.38 | 80.07 |
| SVM | 69.51 | 73.57 | 76.01 | 76.78 | 77.96 | 80.03 | 81.18 |

It can be seen from Table 2 that for the training dataset S. *cerevisiae*, the base-classifiers LightGBM and SVM have different prediction accuracy values for different dimensional reduction methods, and for the same dimensional reduction method, the difference between the prediction accuracy corresponding to the base-classifiers is very small. Among them, LLE has the worst dimensional reduction effect, and the prediction accuracy values of base-classifiers are 69.01% and 69.51%. It shows that this method not only filters out noise information but also removes high-quality classification features at the same time. Compared with the six single feature extraction methods, the dimensional reduction methods MRMD, SE, SVD, and MI can retain the features that are important for model classification and improve the prediction accuracy of the model to a certain extent. However, the prediction accuracy of the four dimensional-reduction methods is still lower than the corresponding value of the original feature dataset All. Compared with the other six dimensional-reduction methods, Elastic Net has the best dimensional reduction effect. The prediction accuracy values of the base-classifiers LightGBM and SVM reach 80.07% and 81.18%, respectively, which are 0.69% and 1.15% higher than the corresponding prediction accuracy value of ET. Considering the calculation speed and prediction performance of the model, we choose Elastic Net as the optimal dimensional reduction method to filter the features that are not relevant and important to the model classification and retain the features that contribute to the prediction model. The Elastic Net improves the prediction performance of the model and effectively distinguishes the real m^6^A sites from non-m^6^A sites.

### 3.5 Predictive performance of StackRAM on the training dataset

Different classifiers have their own unique competitive and prediction advantages and combining all the classifiers does not improve the prediction performance of the model. In the first-stage of the stacked ensemble algorithm, we explore and analyze 7 types of classifiers prediction performance on the training dataset S. *cerevisiae*, including: AdaBoost [57], ERT [56], KNN [58], XGBoost [59], RF [60], LightGBM [61], and SVM [62]. AdaBoost continuously adjusts the training dataset, the learning rate is set to 0.1, and combines the weak learners to obtain a strong classifier. ERT classifier builds 1000 decision random trees and uses Gini index to split nodes. KNN completes the sites recognition task by learning the features of the 50 training samples closest to it. Random forest randomly selects samples and features to avoid the model overfitting, and the number of decision trees is set to 1000. XGBoost sets the learning rate to 0.01 and the tree maximum depth to 10. LightGBM is the tree-based gradient boosting algorithm with a maximum depth of 15 to identify m^6^A sites in the RNA sequences. The support vector machine maps the original feature space to the high-dimensional space through the radial basis kernel function to identify the N^6^-methyladenosine sites. By inputting the optimal feature subset corresponding to Elastic Net into the above 7 classifiers, the performance comparison of 7 machine learning methods on the training dataset S. *cerevisiae* is shown in Fig. 4, and prediction results of 7 machine learning methods on the training dataset S. *cerevisiae* are shown in Supplementary Table S5.

**Fig. 4.**
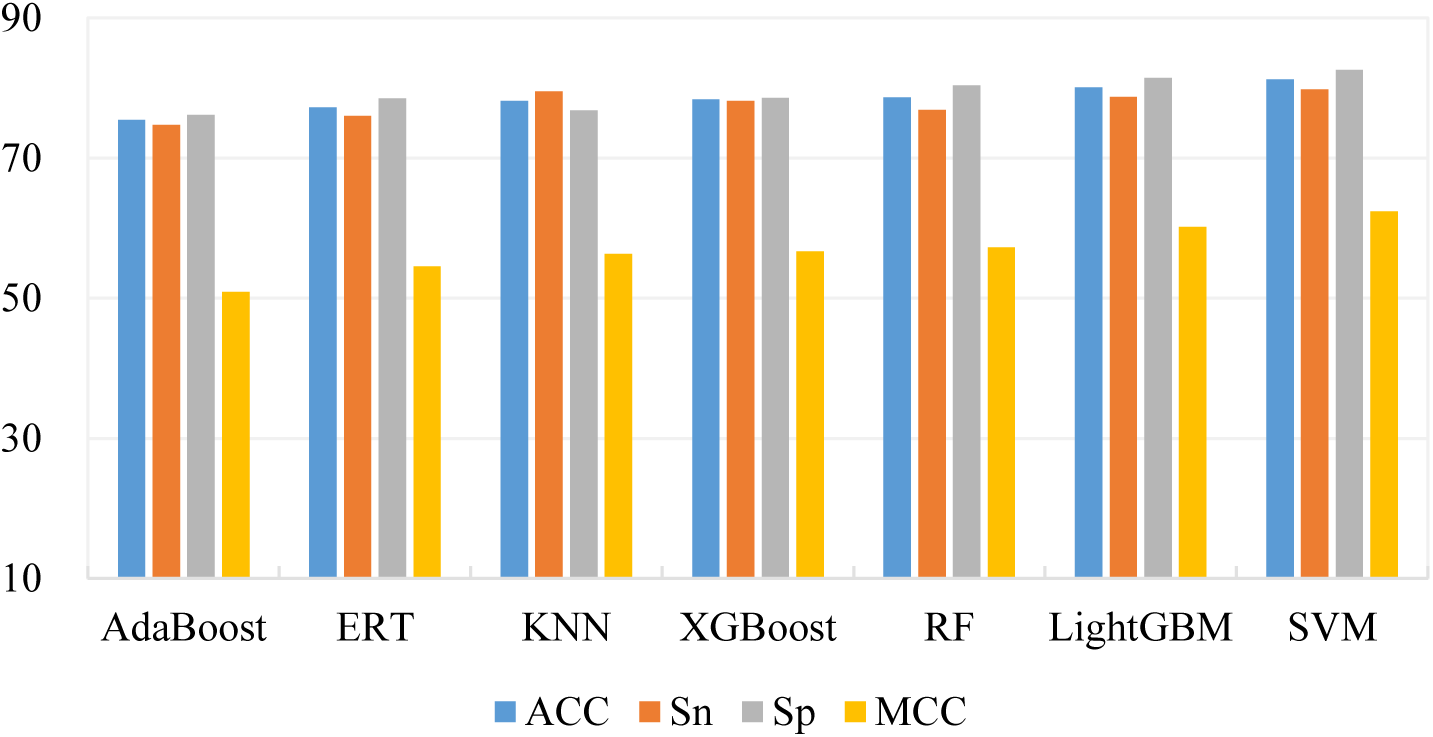
Performance comparison of different classifiers on the training dataset S. *cerevisiae*.

It can be seen intuitively from Fig. 4, different classifiers have different prediction effects on the training dataset S. *cerevisiae*. The prediction accuracy of ERT, KNN, XGBoost, and RF are higher than AdaBoost, but all are lower than 80%. The sensitivity of the classifiers ERT, XGBoost, and RF is lower than the specificity, while the sensitivity of the classifier KNN is higher than the specificity. The prediction accuracy of AdaBoost on the training dataset is 75.44%, the Sn is 74.75%, the Sp is 76.13%, the MCC value is 0.5088, and it has the lowest N^6^-methyladenosine sites recognition ability (Supplementary Table S5). The prediction accuracy of the classifiers LightGBM and SVM are both higher than 80%, reaching 80.07% and 81.18%, respectively. MCC values are also higher than the other five classifiers, which are 0.6016 and 0.6238 (Supplementary Table S5). According to the above prediction results, the LightGBM and SVM are selected as the base-learners of the StackRAM prediction model, which are repeated once. Compared with other stacked ensemble methods, the probability output values of the first stage are directly used as the input feature of the second stage, we combine the probability output values in the first stage and optimal feature subset as a new feature input in the second stage. The combined features are input to the classifiers LR and SVM respectively, and the prediction results of the combined features of the training dataset S. *cerevisiae* of the classifiers LR and SVM are shown in Table 3.

**Table 3.** Prediction results of classifiers LR and SVM on the combined features of training dataset S. *cerevisiae*.

| Classifier | ACC (%) | Sn (%) | Sp (%) | MCC | AUC |
| --- | --- | --- | --- | --- | --- |
| LR | 82.10 | 81.48 | 82.71 | 0.6420 | 0.9011 |
| SVM | 82.56 | 81.94 | 83.17 | 0.6512 | 0.9021 |

According to Table 3, the classifier LR as a classifier in the second stage of the stacked ensemble algorithm, which shows excellent prediction performance. Prediction accuracy value is 82.10%, MCC value is 0.6420, and AUC value is 0.9011. The prediction performance of the classifier SVM is slightly higher than that of LR, and the prediction accuracy value is 82.56%, the MCC value is 0.6512, and the AUC value is 0.9021, which are 0.46%, 0.92%, and 0.1% higher than LR, respectively. SVM combines the combined features to obtain higher prediction results, so SVM is selected as the meta-classifier for the second stage of the StackRAM. In addition, using the above metrics to evaluate the robustness of the model, this paper draws the ROC and PR comparison curves of StackRAM and other classifiers further verify the prediction performance, as shown in Fig. 5.

**Fig. 5.**
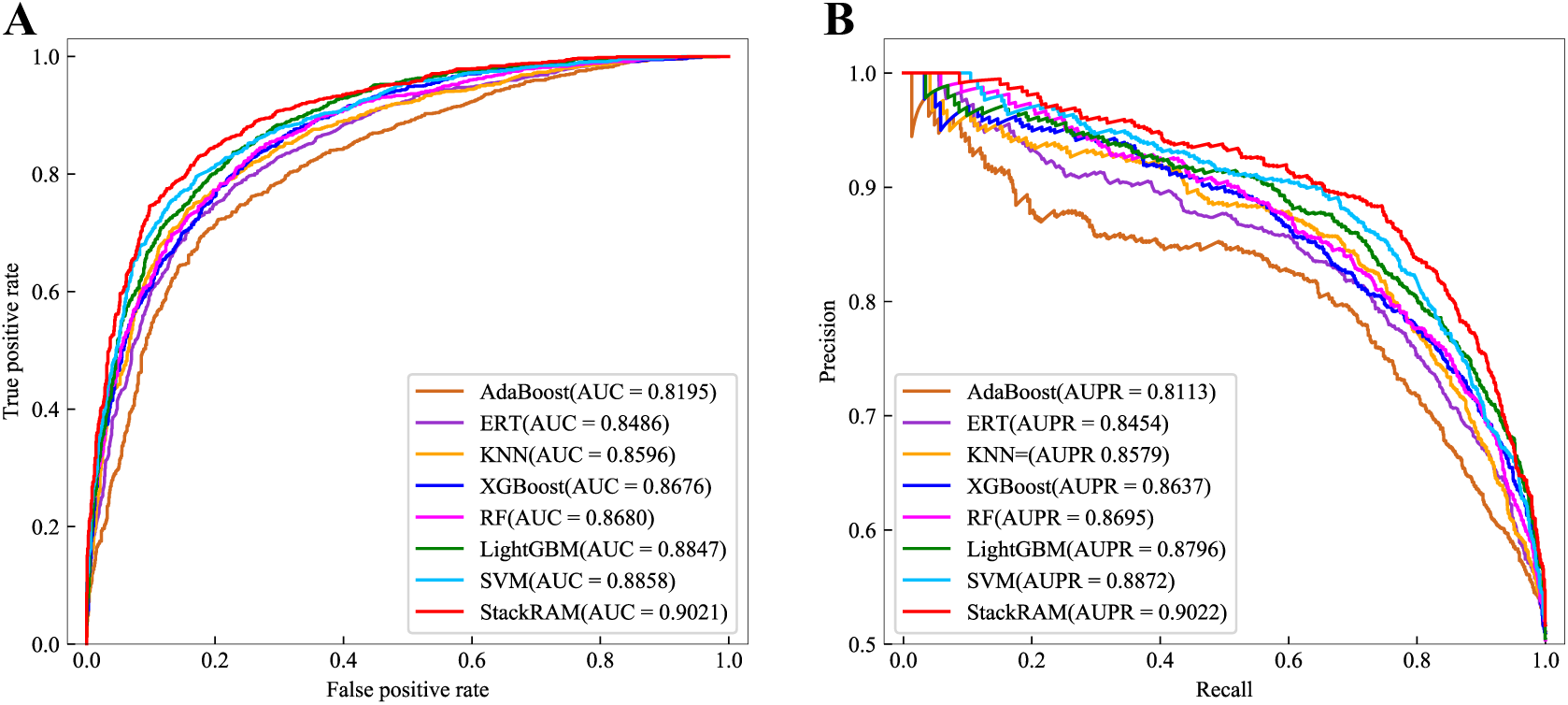
Comparison of ROC and PR curves of StackRAM and other classifiers on the training dataset S. *cerevisiae*.

It can be seen intuitively according to Fig. 5, that the ROC and PR curves of StackRAM on the training dataset S. *cerevisiae* include the curves corresponding to the other 7 classifiers, indicating that the ensemble algorithm can improve the robustness of the model. By comparing the AUC and AUPR values of the corresponding areas under the ROC and PR curves, we find that the AUC value of StackRAM is 0.9021, which is 8.26%5.35%4.25%3.45%3.41%1.74% and 1.63% higher than AdaBoost, ERT, KNN, XGBoost, RF, LightGBM, and SVM, respectively. The AUPR value of StackRAM is 0.9022, which is 9.09%, 5.68%, 4.43%, 3.85%, 3.27%, 2.26% and 1.5% higher han AdaBoost, ERT, KNN, XGBoost, RF, LightGBM and SVM, respectively. Compared with other classifiers, StackRAM integrates single classifier to obtain combined learner with high generalization performance, learns the relationship between different predictors and real classes, and effectively mines sequences that characterize m^6^A sites in RNA sequences.

### 3.6 Comparison with other state-of-the-art methods

To validate the effectiveness of the proposed method, we chose four other state-of-the-art methods to identify m^6^A sites for the dataset S. *cerevisiae*, including iRNA-Methyl [30], pRNAm-PC [63], RNA-Methylpred [64], and Deepm^6^Apred [65]. iRNA-Methyl used pseudo nucleotide composition to identify N^6^-methyladenosine sites. pRNAm-PC predicted the N^6^-methyladenosine sites in the RNA sequences based on physical-chemical properties. RNA-Methylpred incorporated bi-profile Bayes, dinucleotide composition, and k nearest neighbor (KNN) scores for identifying m^6^A sites in RNA. Deepm^6^Apred improved the prediction of N^6^-methyladenosine sites by integrating deep feature representations and handcrafted features. We obtain StackRAM and four prediction methods prediction results on S. *cerevisiae* as shown in Table 4.

**Table 4.**
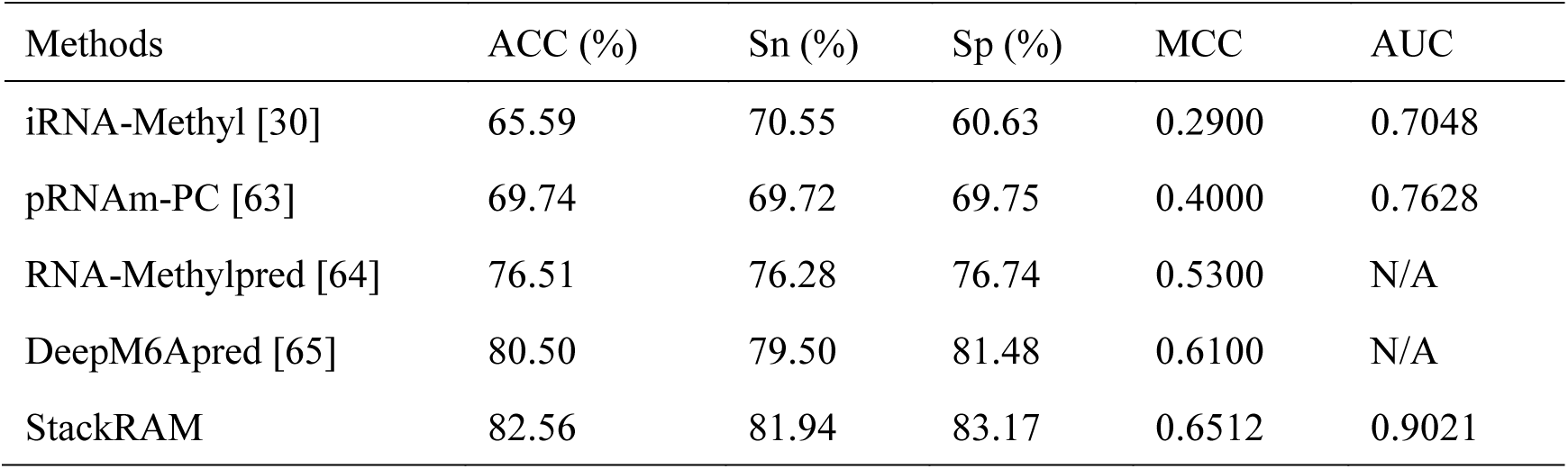
The results comparison of different prediction methods on the training dataset S. *cerevisiae*.

For a fair comparison, the above five prediction methods all use the jackknife method to evaluate the dataset S. *cerevisiae* and then obtain the same evaluation metrics. From Table 4, for the existing prediction methods, Deepm^6^Apred has a better prediction result for the m^6^A sites in RNA. The ACC, Sn, Sp, and MCC values are 80.50%, 79.50%, 81.48%, and 0.61, respectively. However, the novel method StackRAM proposed in this paper has better recognition performance than Deepm^6^Apred. The ACC, Sn, Sp, and MCC values are 2.06%, 2.44%, 1.69%, 4.12% higher than Deepm^6^Apred, respectively. The ACC values are 16.97%, 12.82%, and 6.05% higher than the prediction methods iRNA-Methyl, PRNAm-PC, and RNA-Methylpred, respectively.

The AUC values are 19.73% and 13.93% higher than the prediction methods iRNA-Methyl and PRNAm-PC, respectively. In order to test the generalization ability and robustness of StackRAM, H. *sapiens* and A. *thaliana* are selected as independent test datasets to evaluate the method proposed in this paper. The results of StackRAM and other prediction methods on independent test datasets are shown in Table 5.

**Table 5.** The results comparison of the StackRAM and other methods on independent test datasets.

| Species | Methods | ACC (%) | Sn (%) | Sp (%) | MCC | AUC |
| --- | --- | --- | --- | --- | --- | --- |
| *H. sapiens* | Feng's method [31] | 90.38 | 81.86 | 99.11 | 0.8200 | 0.8490 |
|  | StackRAM | 92.30 | 87.70 | 96.90 | 0.8496 | 0.9617 |
| *A. thaliana* | Chen's method [32] | 84.39 | 68.78 | 100 | 0.7200 | 0.8462 |
|  | StackRAM | 87.06 | 83.76 | 90.36 | 0.7427 | 0.9533 |

As can be seen from Table 5, compared to the other methods prediction results in independent test datasets, StackRAM has the advantage of identifying m^6^A sites. StackRAM prediction accuracy, MCC value, AUC value for H. *sapiens* are 92.30%, 0.8496, 0.9617, which are 1.92%, 2.96%, and 11.27% higher than Feng’s method, respectively. For dataset A. *thaliana*, although the Sp of Chen’s method reaches 100%, the ACC, Sn, MCC and AUC values are 2.67%, 14.98%, 2.27%, and 10.71% lower than StackRAM, respectively. In summary, these results further validate the effectiveness and robustness of StackRAM, indicating that StackRAM is a powerful prediction method that is not only competitive for training dataset m^6^A sites recognition but also has better prediction performance in cross-species sites recognition.

## 4. Conclusion

This paper proposes an accurate and robust cross-species calculational method StackRAM, which predicts the m^6^A sites based on RNA nucleotide sequence information. Firstly, the RNA sequence is encoded by six sequence-derived feature extraction methods. Compared with the single feature extraction method, multi-information fusion can extract features from sequence information and physicochemical properties, which fully reflects the characteristics difference between m^6^A sites and non-m^6^A sites. Secondly, Elastic Net is used to optimize the original feature space for the first time. Compared with other dimensional reduction methods, the Elastic Net can retain important features related to model classification and filter redundant and noise information. Then, by comparing the prediction results of multiple classifiers on the optimal feature subset, LightgGBM and SVM are selected as the base-classifiers of the first stage of StackRAM, and SVM is used as the second-stage meta-classifier. In addition, the StackRAM method combines the output probability of the first stage with the optimal feature subset, and the combined feature as the input feature of the second stage to improve the m^6^A sites prediction performance of the model. Compared with other state-of-the-art prediction methods, the novel method proposed in this paper has a higher prediction success rate on various evaluation metrics, indicating that the ensemble method can use different base-classifiers to predict the m^6^A sites through hierarchical learning. Finally, the evaluation results of the independent test datasets show that StackRAM can effectively capture the necessary characteristics of RNA methylation modification sites and has the robustness to identify m^6^A sites in different species. The StackRAM approach not only improves understanding of epigenetic modification of different biological processes but also impacts drug development. Although the method proposed in this paper can quite accurately predict the N^6^-methyladenosine sites of RNA, and there is still room for further improvement. In the future, post-translational modification sites datasets of different species and types will be generated, and we will integrate deep learning methods and traditional machine learning methods to train StackRAM to detect more RNA post-translational modifications via integrating more feature extraction methods.

## Declaration of competing interest

No author associated with this paper has disclosed any potential or pertinent conflicts which may be perceived to have impending conflict with this work.

## Acknowledgments

This work was supported by the National Nature Science Foundation of China (No. 61863010), the Key Research and Development Program of Shandong Province of China (No. 2019GGX101001), and the Natural Science Foundation of Shandong Province of China (No. ZR2018MC007, ZR2019MEE066).

## Notes

### Competing Interest Statement

The authors have declared no competing interest.

## References

[1] I.A. Roundtree, M.E. Evans, T. Pan, C. He, Dynamic RNA modifications in gene expression regulation, Cell 169 (2017) 1187–1200.

[2] M.A. Machnicka, K. Milanowska, O.O. Oglou, E. Purta, M. Kurkowska, A. Olchowik, W. Januszewski, S. Kalinowski, S. Dunin-Horkawicz, K.M. Rother, M. Helm, J.M. Bujnicki, H. Grosjean, MODOMICS: a database of RNA modification pathways—2013 update, Nucleic Acids Res. 41 (2013) D262–D267.

[3] Y. Wan, K. Tang, D. Zhang, S. Xie, X. Zhu, Z. Wang, Z. Lang, Transcriptome-wide high-throughput deep m^6^A-seq reveals unique differential m^6^A methylation patterns between three organs in Arabidopsis thaliana, Genome Biol. 16 (2015) 272.

[4] W. Chen, H. Tran, Z. Liang, H. Lin, L. Zhang, Identification and analysis of the N^6^-methyladenosine in the Saccharomyces cerevisiae transcriptome, Sci. Rep. 5 (2015) 13859.

[5] X. Deng, K. Chen, G.Z. Luo, X. Weng, Q. Ji, T. Zhou, C. He, Widespread occurrence of N^6^-methyladenosine in bacterial mRNA, Nucleic Acids Res. 43 (2015) 6557–6567.

[6] W. Huang, J. Xiong, Y. Yang, S.M. Liu, B.F. Yuan, Y.Q. Feng, Determination of DNA adenine methylation in genomes of mammals and plants by liquid chromatography/mass spectrometry, Rsc Adv. 5 (2015) 64046–64054.

[7] X. Wang, Z. Lu, A. Gomez, G.C. Hon, Y. Yue, D. Han, Y. Fu, M. Parisien, Q. Dai, G. Jia, B. Ren, T. Pan, C. He, N^6^-methyladenosine-dependent regulation of messenger RNA stability, Nature 505 (2014) 117–120.

[8] N. Liu, Q. Dai, G. Zheng, C. He, M. Parisien, T. Pan, N^6^-methyladenosine-dependent RNA structural switches regulate RNA-protein interactions, Nature 518 (2015) 560–564.

[9] Y. Wang, Y. Li, J.I. Toth, M.D. Petroski, Z. Zhang, J.C. Zhao, N^6^-methyladenosine modification destabilizes developmental regulators in embryonic stem cells, Nat. Cell Biol. 16 (2014) 191–198.

[10] Y. Yang, B.F. Sun, W. Xiao, X. Yang, H.Y. Sun, Y.L. Zhao, Y.G. Yang, Dynamic m^6^A modification and its emerging regulatory role in mRNA splicing, Sci. Bull. 60 (2015) 21–32.

[11] Y. Niu, X. Zhao, Y.S. Wu, M.M. Li, X.J. Wang, Y.G. Yang, N^6^-methyl-adenosine (m^6^A) in RNA: an old modification with a novel epigenetic function, Genom. Proteom. Bioinf. 11 (2013) 8–17.

[12] C. Zhang, D. Samanta, H. Lu, J.W. Bullen, H. Zhang, I. Chen, X. He, G.L. Semenza, Hypoxia induces the breast cancer stem cell phenotype by HIF-dependent and ALKBH5-mediated m^6^A-demethylation of NANOG mRNA, P. Natl. Acad. Sci. 113 (2016) E2047–E2056.

[13] M. Brocard, A. Ruggieri, N. Locker, m^6^A RNA methylation, a new hallmark in virus-host interactions, J. Gen. Virol. 98 (2017) 2207–2214.

[14] K.D. Meyer, Y. Saletore, P. Zumbo, O. Elemento, C.E. Mason, S.R. Jaffrey, Comprehensive analysis of mRNA methylation reveals enrichment in 3′ UTRs and near stop codons, Cell 149 (2012) 1635–1646.

[15] G. Keith, Mobilities of modified ribonucleotides on two-dimensional cellulose thin-layer chromatography, Biochimie 77 (1995) 142–144

[16] G. Zheng, J.A. Dahl, Y. Niu, P. Fedorcsak, C.M. Huang, C.J. Li, C.B. Vågbø, Y. Shi, W.L. Wang, S.H. Song, Z. Lu, R.P.G. Bosmans, Q. Dai, Y.J. Hao, X. Yang, W.M. Zhao, W.M. Tong, X.J. Wang, F. Bogdan, K. Furu, Y. Fu, G. Jia, X. Zhao, J. Liu, H.E. Krokan, A. Klungland, Y.G. Yang, C. He, ALKBH5 is a mammalian RNA demethylase that impacts RNA metabolism and mouse fertility, Mol. Cell 49 (2013) 18–29.

[17] D. Dominissini, S. Moshitch-Moshkovitz, M. Salmon-Divon, N. Amariglio, G. Rechavi, Transcriptome-wide mapping of N^6^-methyladenosine by m^6^A-seq based on immunocapturing and massively parallel sequencing, Nat. Protoc. 8 (2013) 176.

[18] Y. Zhou, P. Zeng, Y.H. Li, Z. Zhang, Q. Cui, SRAMP: prediction of mammalian N^6^-methyladenosine (m^6^A) sites based on sequence-derived features, Nucleic Acids Res. 44 (2016) e91.

[19] Y. Huang, N. He, Y. Chen, Z. Chen, L. Li, BERMP: a cross-species classifier for predicting m^6^A sites by integrating a deep learning algorithm and a random forest approach, Int. J. Biol. Sci. 14 (2018) 1669–1677.

[20] Z. Zhao, H. Peng, C. Lan, Y. Zheng, L. Fang, J. Li, Imbalance learning for the prediction of N^6^-Methylation sites in mRNAs, BMC Genomics 19 (2018) 574.

[21] W. Chen, H. Ding, X. Zhou, H. Lin, K.C. Chou, iRNA(m^6^A)-PseDNC: identifying N^6^-methyladenosine sites using pseudo dinucleotide composition, Anal. Biochem. 561 (2018) 59–65.

[22] W. Chen, P. Xing, Q. Zou, Detecting N^6^-methyladenosine sites from RNA transcriptomes using ensemble Support Vector Machines, Sci. Rep. 7 (2017) 40242.

[23] X. Wang, R. Yan, RFAthM6A: a new tool for predicting m^6^A sites in Arabidopsis thaliana, Plant Mol. Boil. 96 (2018) 327–337.

[24] S. Akbar, M. Hayat, iMethyl-STTNC: identification of N^6^-methyladenosine sites by extending the idea of SAAC into Chou′s PseAAC to formulate RNA sequences, J. Theor. Biol. 455 (2018) 205–211.

[25] J. Zhang, P. Feng, H. Lin, W. Chen, Identifying RNA N^6^-methyladenosine sites in escherichia coli genome, Front. Microbiol. 9 (2018) 955.

[26] M. Zhang, J.W. Sun, Z. Liu, M.W. Ren, H.B. Shen, D.J. Yu, Improving N^6^-methyladenosine site prediction with heuristic selection of nucleotide physical-chemical properties, Anal. Biochem. 508 (2016) 104–113.

[27] S. Xiang, K. Liu, Z. Yan, Y. Zhang, Z. Sun, RNAMethPre: a web server for the prediction and query of mRNA m^6^A sites, PLoS One 11 (2016) e0162707.

[28] X. Qiang, H. Chen, X. Ye, R. Su, L. Wei, M6AMRFS: robust prediction of N6-methyladenosine sites with sequence-based features in multiple species, Front. Genet. 9 (2018) 495.

[29] W. Chen, H. Tang, H. Lin, MethyRNA: a web server for identification of N^6^-methyladenosine sites, J. Biomol. Struct. Dyn. 35 (2017) 683–687.

[30] W. Chen, P. Feng, H. Ding, H. Lin, K.C. Chou, iRNA-Methyl: Identifying N^6^-methyladenosine sites using pseudo nucleotide composition, Anal. Biochem. 490 (2015) 26–33.

[31] P. Feng, H. Ding, H. Yang, W. Chen, H. Lin, K.C. Chou, iRNA-PseColl: identifying the occurrence sites of different RNA modifications by incorporating collective effects of nucleotides into PseKNC, Mol. Ther.-Nucl. Acids 7 (2017) 155–163.

[32] W. Chen, P. Feng, H. Ding, H. Lin, Identifying N^6^-methyladenosine sites in the Arabidopsis thaliana transcriptome, Mol. Genet. Genomics 291 (2016) 2225–2229.

[33] S. Xiang, Z. Yan, K. Liu, Y. Zhang, Z. Sun, AthMethPre: A web server for the prediction and query of mRNA m^6^A sites in Arabidopsis thaliana, Mol. BioSyst. 12 (2016) 3333–3337.

[34] W. Chen, P. Feng, H. Yang, H. Ding, H. Lin, K.C. Chou, iRNA-3typeA: identifying three types of modification at RNA′s adenosine sites, Mol. Ther.-Nucl. Acids 11 (2018) 468–474.

[35] W. Chen, P. Feng, H. Tang, H. Ding, H. Lin, Identifying 2′-O-methylationation sites by integrating nucleotide chemical properties and nucleotide compositions, Genomics 107 (2016) 255–258.

[36] G.Q. Li, Z. Liu, H.B. Shen, D.J. Yu, TargetM6A: Identifying N^6^-Methyladenosine Sites From RNA Sequences via Position-Specific Nucleotide Propensities and a Support Vector Machine, IEEE T. Nanobiosc. 15 (2016) 674–682.

[37] W. Chen, T.Y. Lei, D.C. Jin, H. Lin, K.C. Chou, PseKNC: a flexible web server for generating pseudo K-tuple nucleotide composition, Anal. Biochem. 456 (2014) 53–60.

[38] Z. Chen, P. Zhao, F. Li, T.T. Marquez-Lago, A. Leier, J. Revote, Y. Zhu, D.R. Powell, T. Akutsu, G.I. Webb, K.C. Chou, A.I. Smith, R.J. Daly, J. Li, J. Song, iLearn: an integrated platform and meta-learner for feature engineering, machine-learning analysis and modeling of DNA, RNA and protein sequence data, Brief. Bioinform. (2019) https://doi.org/10.1093/bib/bbz041.

[39] R. Tibshirani, Regression shrinkage and selection via the lasso, J. R. Stat. Soc. B. 58 (1996) 267–288.

[40] H. Zou, T. Hastie, Regularization and variable selection via the elastic net, J. R. Stat. Soc. B. 67 (2005) 301–320.

[41] S. Saha, S. Mitra, R.K. Yadav, A stack-based ensemble framework for detecting cancer microRNA biomarkers, Genom. Proteom. Bioinf. 15 (2017) 381–388.

[42] A. Mishra, P. Pokhrel, M.T. Hoque, StackDPPred: a stacking based prediction of DNA-binding protein from sequence, Bioinformatics 35 (2019) 433–441.

[43] Y. Xiong, Q. Wang, J. Yang, X. Zhu, D. Wei, PredT4SE-stack: prediction of bacterial type IV secreted effectors from protein sequences using a stacked ensemble method, Front. Microbiol. 9 (2018) 2571.

[44] R. Su, X. Liu, G. Xiao, L. Wei, Meta-GDBP: a high-level stacked regression model to improve anticancer drug response prediction, Brief. Bioinform. (2019) https://doi.org/10.1093/bib/bbz022.

[45] C. Chen, Q. Zhang, Q. Ma, B. Yu, LightGBM-PPI: Predicting protein-protein interactions through LightGBM with multi-information fusion, Chemometr. Intell. Lab. 191 (2019) 54–64.

[46] H. Shi, S. Liu, J. Chen, X. Li, Q. Ma, B. Yu, Predicting drug-target interactions using Lasso with random forest based on evolutionary information and chemical structure, Genomics 111 (2019) 1839–1852.

[47] B. Yu, S. Li, W. Qiu, M. Wang, J. Du, Y. Zhang, X. Chen, Prediction of subcellular location of apoptosis proteins by incorporating PsePSSM and DCCA coefficient based on LFDA dimensionality reduction, BMC Genomics 19 (2018) 478.

[48] H. Zhou, C. Chen, M. Wang, Q. Ma, B. Yu, Predicting Golgi-Resident Protein Types Using Conditional Covariance Minimization With XGBoost Based on Multiple Features Fusion, IEEE Access 7 (2019) 144154–144164.

[49] X. Wang, B. Yu, A. Ma, C. Chen, B. Liu, Q. Ma, Protein-protein interaction sites prediction by ensemble random forests with synthetic minority oversampling technique, Bioinformatics 35 (2019) 2395–2402.

[50] X. Cui, Z. Yu, B. Yu, M. Wang, B. Tian, Q. Ma, UbiSitePred: A novel method for improving the accuracy of ubiquitination sites prediction by using LASSO to select the optimal Chou’s pseudo components, Chemometr. Intell. Lab.184 (2019) 28–43.

[51] S.T. Roweis, L.K. Saul, Nonlinear dimensionality reduction by locally linear embedding, Science 290 (2000) 2323–2326.

[52] Q. Zou, J. Zeng, L. Cao, R. Ji, A novel features ranking metric with application to scalable visual and bioinformatics data classification, Neurocomputing 173 (2016) 346–354.

[53] A.Y. Ng, M.I. Jordan, Y. Weiss, On spectral clustering: Analysis and an algorithm, in: Advances in Neural Information Processing Systems, 2002, pp. 849–856.

[54] M.E. Wall, A. Rechtsteiner, L.M. Rocha, Singular value decomposition and principal component analysis, in: A Practical Approach to Microarray Data Analysis, 2003, pp. 91–109.

[55] B.C. Ross, Mutual information between discrete and continuous data sets, PLoS One 9 (2014) e87357.

[56] P. Geurts, D. Ernst, L. Wehenkel, Extremely randomized trees, Mach. Learn. 63 (2006) 3–42.

[57] Y. Freund, R.E. Schapire, A decision-theoretic generalization of on-line learning and an application to boosting, J. Comput. Syst. Sci. 55 (1997) 119–139.

[58] F. Nigsch, A. Bender, B. van Buuren, J. Tissen, E. Nigsch, J.B.O. Mitchell, Melting point prediction employing k-nearest neighbor algorithms and genetic parameter optimization, J. Chemical. Inf. Model. 46 (2006) 2412–2422.

[59] T. Chen, C. Guestrin, Xgboost: A scalable tree boosting system, in: Proceedings of the 22nd ACM SIGKDD International Conference On Knowledge Discovery And Data Mining, 2016, pp. 785–794.

[60] L. Breiman, Random forest, Mach. Learn. 45 (2001) 5–32.

[61] G. Ke, Q. Meng, T. Finley, T. Wang, W. Chen, W. Ma, W. Ma, Q. Ye, T.Y. Liu, Lightgbm: A highly efficient gradient boosting decision tree, in: Advances in Neural Information Processing Systems, 2017, pp. 3146–3154.

[62] C. Cortes, V. Vapnik, Support-vector networks, Mach. Learn. 20 (1995) 273–297.

[63] Z. Liu, X. Xiao, D.J. Yu, J. Jia, W.R. Qiu, K.C. Chou, pRNAm-PC: Predicting N^6^-methyladenosine sites in RNA sequences via physical-chemical properties, Anal. Biochem. 497 (2016) 60–67.

[64] C.Z. Jia, J.J. Zhang, W.Z. Gu, RNA-MethylPred: a high-accuracy predictor to identify N6-methyladenosine in RNA, Anal. Biochem. 510 (2016) 72–75.

[65] L. Wei, R. Su, B. Wang, X. Li, Q. Zou, X. Gao, Integration of deep feature representations and handcrafted features to improve the prediction of N^6^-methyladenosine sites, Neurocomputing 324 (2019) 3–9.

