## Supplementary Tables for "StackRAM: a cross-species method for identifying RNA N^6^-methyladenosine sites based on stacked ensemble"

**Table S1.** Prediction accuracy values of S. *cerevisiae* for different parameters $\lambda$ in PseDNC.

**Table S2.** Dimensions of S. *cerevisiae* for different feature extraction methods.

**Table S3.** Elastic Net dimensions and base-classifiers ACC values for different $\lambda_2$ when $\lambda_1 = 0.1$.

**Table S4.** Elastic Net dimensions and base-classifiers ACC values for different $\lambda_1$ when $\lambda_2 = 0.05$.

**Table S5.** Jackknife method prediction results of S. *cerevisiae* for different classifiers.

**Table S1.**

Prediction accuracy values of S. *cerevisiae* for different parameters $\lambda$ in PseDNC.

| λ | 4 | 8 | 12 | 16 | 20 | 23 |
|---|---|---|---|---|---|---|
| LightGBM | 60.37 | 62.13 | 62.70 | 62.01 | 61.82 | 63.16 |
| SVM | 62.70 | 64.54 | 64.23 | 64.54 | 64.08 | 64.38 |

**Table S2.**

Dimensions of S. cerevisiae for different feature extraction methods.

| Methods | ANF | Binary | NCP | K-mer | PseDNC | PSTNP | All |
|---|---|---|---|---|---|---|---|
| dimensions | 50 | 204 | 153 | 16 | 39 | 49 | 511 |

**Table S3.**

Elastic Net dimensions and base-classifiers ACC values for different $\lambda_2$ when $\lambda_1 = 0.1$.

| $\lambda_2$ | 0.01 | 0.03 | 0.05 | 0.07 | 0.09 |
|---|---|---|---|---|---|
| dimensions | 249 | 200 | 165 | 136 | 101 |
| LightGBM | 78.92 | 79.27 | 80.07 | 79.38 | 79.34 |
| SVM | 80.60 | 79.57 | 81.18 | 80.06 | 81.10 |

**Table S4.**

Elastic Net dimensions and base-classifiers ACC values for different $\lambda_1$ when $\lambda_2 = 0.05$.

| $\lambda_1$ | 0.1 | 0.3 | 0.5 | 0.7 | 0.9 |
|---|---|---|---|---|---|
| dimensions | 165 | 68 | 63 | 63 | 61 |
| LightGBM | 80.07 | 78.81 | 79.30 | 79.30 | 61.21 |
| SVM | 81.18 | 72.61 | 65.26 | 66.41 | 61.21 |

Elastic Net adjusts the convex combination between $\ell_1$ and $\ell_2$ by penalty parameter $\lambda_2$, and then selects features with correlation. If the penalty parameters $\lambda_1$ and $\lambda_2$ are set too large at the same time, all features will be eliminated. Therefore, this paper first fixed the penalty parameter $\lambda_1$ to 0.1, and set the parameters $\lambda_2$ to 0.01, 0.03, 0.05, 0.07, 0.09 in order. Determine the value of the parameter $\lambda_2$ as 0.05 based on the prediction accuracy of the base-classifiers LightGBM and SVM. Then set the parameter values of $\lambda_1$ to 0.1, 0.3, 0.5, 0.7, and 0.9 in order, the value of parameter $\lambda_1 = 0.1$ according to the prediction accuracy of the base-classifiers.

It can be seen intuitively from Table S3, as the penalty parameters increase, the corresponding feature subset dimensions change in the opposite direction. For the base-classifier LightGBM, the prediction accuracy of different feature subsets first increases and then decreases. For the base-classifier SVM, the prediction accuracy of different feature subsets fluctuates greatly.

When $\lambda_2$ is 0.05, the dimensions of the corresponding optimal feature subset is 165, and the base-classifiers have the highest prediction accuracy, 80.07% and 81.18%, respectively. It can be seen intuitively from Table S4, for the base-classifier LightGBM, as the value of the parameter $\lambda_1$ increases, the prediction accuracy of different feature subsets shows a downward trend. For the base-classifier SVM, when the parameter $\lambda_1 = 0.1$, it has the highest prediction accuracy. Considering the prediction accuracy values of the base-classifiers on the jackknife method, we determine that the optimal parameters of Elastic Net $\lambda_1$ is 0.1 and $\lambda_2$ is 0.05.

**Table S5.**

Jackknife method prediction results of different classifiers for S. *cerevisiae.*

| Classifiers | ACC (%) | Sn (%) | Sp (%) | MCC | AUC |
|---|---|---|---|---|---|
| AdaBoost | 75.44 | 74.75 | 76.13 | 0.5088 | 0.8195 |
| ERT | 77.24 | 75.98 | 78.50 | 0.5449 | 0.8486 |
| KNN | 78.16 | 79.50 | 76.82 | 0.5633 | 0.8596 |
| XGBoost | 78.35 | 78.12 | 78.58 | 0.5670 | 0.8676 |
| RF | 78.62 | 76.89 | 80.34 | 0.5726 | 0.8680 |
| LightGBM | 80.07 | 78.73 | 81.41 | 0.6016 | 0.8847 |
| SVM | 81.18 | 79.80 | 82.56 | 0.6238 | 0.8858 |
